# MicroRNA-148a Regulates Low-Density Lipoprotein Metabolism by Repressing the (Pro)renin Receptor

**DOI:** 10.1101/831529

**Authors:** Na Wang, Lishu He, Hui Lin, Lunbo Tan, Yuan Sun, Xiaoying Zhang, A.H. Jan Danser, Hong S. Lu, Yongcheng He, Xifeng Lu

## Abstract

High plasma LDL cholesterol (LDL-c) concentration is a major risk factor for atherosclerosis. Hepatic LDLR regulates LDL metabolism, and thereby plasma LDL-c concentration. Recently, we identified the (pro)renin receptor [(P)RR] as a novel regulator of LDL metabolism, which regulates LDLR degradation and hence its protein abundance and activity. *In silicon* analysis suggests that the (P)RR is a target of miR-148a. In this study we determined whether miR-148a could regulate LDL metabolism by regulating *(P)RR* expression in HepG2 and Huh7 cells. We found that miR-148a suppressed *(P)RR* expression by binding to the 3’-untranslated regions (3’-UTR) of (P)RR mRNA. Mutating the binding sites for miR-148a in the 3’-UTR of (P)RR mRNA abolished the inhibitory effects of miR-148a on *(P)RR* expression. In line with our recent findings, reduced *(P)RR* expression resulted in decreased cellular LDL uptake, likely as a consequence of decreased LDLR protein abundance. Overexpressing the (P)RR prevented miR-148a-induced reduction in LDLR abundance and cellular LDL uptake. Our study supports a new concept that miR-148a is a regulator of *(P)RR* expression. By reducing (P)RR abundance, miR-148a decreases LDLR protein abundance and consequently cellular LDL uptake.

## Introduction

High plasma LDL cholesterol (LDL-c) level is a major risk factor for CVD, a leading cause of world-wide death. Plasma LDL is mainly cleared by the LDLR in the liver. Genetic mutations resulting in defective LDLR functions are associated with elevated plasma LDL-c levels and increased risks for CVD [1]. Additionally, decreased LDLR protein abundance, caused either by reduced transcription or increased protein degradation, leads to disturbed LDL clearance. Due to its importance in regulating LDL metabolism, hepatic LDLR transcription is tightly controlled under normal physiological state by sterol regulatory element-binding protein in response to cellular cholesterol levels [2]. Protein degradational regulation also controls LDLR abundance. Several factors, such as protein convertase subtilisin/kexin9 (PCSK9) and inducible degrader of LDLR (IDOL), have been found to regulate LDLR protein degradation [3–5]. Dysregulation of these regulatory factors, caused by genetic mutations or pathological conditions, is associated with altered plasma LDL levels and CVD risks [6, 7].

Recently, we have identified the (pro)renin receptor [(P)RR] as a novel regulator of LDLR degradation [8]. The (P)RR can bind renin and prorenin with higher affinity, playing an important role in local activation of the renin-angiotensin system (RAS). Upon renin/prorenin binding, the (P)RR triggers intracellular signaling cascades, such as signal-regulated kinase 1/2 and p38 kinase signaling, resulting in upregulation of profibrotic factors including transforming growth factor β, collagen-1 and fibronectin [9–11]. The (P)RR has also been reported to play crucial roles in maintaining vacuolar H^+^-ATPase integrity, Wnt/β-catenin and PCP signaling, glucose metabolism, and lipid metabolism, independent of renin/prorenin [12–16].

Despite its importance in several physiological processes, it is unclear how the (P)RR itself is being regulated. Interestingly, renin/prorenin-(P)RR signaling can reduce (P)RR expression via a negative feedback mechanism [17]. Altered (P)RR expression has been reported in several tissues under pathophysiological conditions, such as in the adipose tissue of HFD-fed obese mice and in the kidney of diabetic nephropathy (DN) mice [18], and may play a role in the onset and development of such diseases.

Recent studies have found that several microRNAs (miRNAs) regulate (P)RR expression, suggesting that miRNAs may play a role in regulating (P)RR functions [19]. MiRNAs are short non-coding RNAs about 22 nucleotides in length, which can affect protein abundance of target genes via non-sense mediated messenger RNA decay or translational inhibition by binding to the 3’-untraslated regions (3’-UTR) of the mRNA of target gene [20]. With their ability to regulate gene expression, miRNAs have been reported to play a role in a wide variety of pathophysiological processes, such as hypertension, obesity, and atherosclerosis [21–23]. Using *in silicon* analysis, we found that (P)RR is a potential target of miR-148a. Moreover, increased circulating miR-148a levels are associated with elevated plasma LDL-c levels, linking miR-148a with plasma cholesterol homeostasis regulation [24]. As we recently discovered, inhibiting the (P)RR reduces LDLR protein abundance and thus affecting LDL metabolism both *in vitro* and *in vivo* [8, 16]. We hypothesized that miR-148a regulates LDL metabolism by down-regulating the (P)RR.

To test this hypothesis, we determined whether miR-148a affects *(P)RR* expression, and its consequences on cellular LDL uptake in human hepatic cells. We found that miR-148a potently reduced *(P)RR* expression and cellular LDL uptake in HepG2 and Huh7 cells. Blocking miR-148a using its inhibitors attenuated miR-148a-induced reduction in *(P)RR* expression and cellular LDL uptake. Furthermore, we found that overexpressing the (P)RR prevented miR-148a-induced reduction in cellular LDL uptake. Our study thus shows that the (P)RR is a key mediator for miR-148a to regulate LDL metabolism.

## Materials and Methods

### Chemicals and Reagents

All chemicals and reagents were purchased from Sigma Aldrich with highest purity unless where else indicated.

### Cell Culture and Transfection

HepG2, Huh7 and HEK293 cells were maintained in DMEM GlutaMAX (Thermo Fisher Scientific) supplemented with 10% fetal bovine serum (Thermo Fisher Scientific), 100 U/mL penicillin and 100 mg/mL streptomycin (Thermo Fisher Scientific) at 37 °C and 5% CO_2_. For miRNA mimics transfection in HEK293, HepG2 and Huh7 cells, Lipofectamine RNAimax (Thermo Fisher Scientific) was used following the manufacturer’s protocol. MiRNA mimics and inhibitors were synthesized by GenePharm (Suzhou, China) and their sequences were listed in **Table S1**. For plasmid transfection, Lipofectamine 2000 (Thermo Fisher Scientific) was used for transfecting HEK293 cells, and Lipofectamine 3000 (Thermo Fisher Scientific) was used for transfecting HepG2 and Huh7 cells. Unless specially indicated otherwise, HepG2 and Huh7 cells were cultured in sterol-depleted medium [(DMEM supplemented with 10% bovine lipid-deficient serum (LPDS), 5 μg/mL simvastatin (Selleck), and 100 μM mevalonic acid (Sigma Aldrich)] as previously described [5].

### Immunoblotting

For immunoblotting, cells were washed twice with ice-cold PBS and lysed at 4 °C in RIPA buffer (containing 50 mmol/L Tris-HCl, 150 mmol/L NaCl, 1% Triton X-100, 0.1% SDS, 1% sodium deoxycholate, and complete protease inhibitor; pH 7.4). Lysates were centrifuged at 10,000 g for 10 minutes at 4 °C, and protein concentrations in the supernatant were measured with a bicinchoninic acid assay kit (Thermo Fisher Scientific). Equal amount of proteins (20 – 30 μg) were resolved by 4 – 20% Bis-Tris gels (GeneScript) and transferred to PVDF membranes using iBlot^®^ 2 Dry Blotting System (Thermo Fisher Scientific). Blots were probed using previously described primary antibodies: anti-LDLR (1:1000; Proteintech), anti-(P)RR (1:1000), and anti-tubulin (1:5000; Proteintech), and HRP-conjugated goat antimouse or goat anti-rabbit antibodies (1:5000; Jackson ImmunoResearch) and detected by ECL.

### RNA Isolation and qPCR

Total RNA was isolated from cells using TRIzol (Thermo Fisher Scientific) and Direct-zol^TM^ RNA MiniPrep kit (ZYMO Research) following the manufacturer’s protocol. One microgram of total RNA was reverse transcribed with Prime Script^TM^ RT Master Mix (TaKaRa). Quantitative PCR assays were performed on a qTOWER apparatus (Analytic Jena) using SYBR Green master mix (TaKaRa). Gene expression was normalized for the expression of 36B4, and expressed as mean ± SEM. Sequences of the primers used in this study are provided in **Table S1**.

### LDL Uptake Assay

Blood was drawn from healthy volunteer, and LDL was prepared by ultracentrifugation as previously described [25]. LDL was then labeled with DyLight-488 (Thermo Fisher Scientific) following manufacturer’s protocol. LDL uptake was measured using DyLight-488-labeled LDL as previously described [26]. Briefly, HepG2 or Huh7 cells were incubated in sterol-depleted medium for 16 hours prior to adding LDL. Cells were incubated with 5 μg/mL DyLight-488-labeled LDL in DMEM containing 0.5% BSA for 3 hours at 37 °C or 4 °C. In these experiments, 100 μg/mL non-labeled LDL was used to correct for non-specific LDL binding. After incubation, cells were washed twice by ice-cold PBS supplemented with 0.5% BSA and then lysed in RIPA buffer. Specific LDL uptake was calculated as the fluorescent intensity differences between 37 °C and 4 °C, after subtracting non-specific association/binding. LDL uptake was determined by quantification of the fluorescence signal using a multi-mode fluorescent microplate reader (Cytation 5, Biotek), and corrected for the protein contents in the same lysates as determined with the BCA assay. To visualize LDL uptake, cells were cultured on coverslips and treated as above mentioned, then fixed with 4% paraformaldehyde, and mounted with Vectorshield containing DAPI (Vector Laboratories). Prepared slides were visualized with Cytation 5 using slide imaging mode with a 40x objective lens.

### 3’-UTR Luciferase Reporter Assay

Full length 3’-UTR of human (P)RR and LDLR were cloned into the luciferase reporter vector (Promega). Mutant 3’-UTR luciferase reporter vectors were generated by PCR using site-directed mutagenesis technique. Correctness of the constructs were then confirmed by sequencing. HEK293 were transfected with constructed vectors, together with miR-148a mimics or control oligos. After 48 hours of transfection, cells were lysed using lysis buffer provided by Dual-Glo Luciferase Assay kit, and lysates were cleared as described above. Luciferase activities were measured using Dual-Glo Luciferase Assay (Promega). In short, *Firefly* luciferase activities in the lysates were measured, and corrected for *Renilla* luciferase activities in the same lysates.

### Statistical Analysis

Data are presented as mean ± SEM. Data sets were first analyzed using D’Agostino-Pearson omnibus test for normality. F test, Browne-Forsythe test or Bartlett’s test was performed for testing if the variances of different data sets are equal. When passing normality and equal variance test, parametric Student’s t-test was performed for comparison of two groups, or One-way ANOVA followed by the Bonferroni correction was performed for comparison of more than two groups. When failed passing normality and equal variance test, non-parametric Student’s t-test with Welch’s correction was performed, or non-parametric One-Way ANOWA followed by Dunn’s correction was performed for comparison of more than two groups. P values of <0.05 were considered significant. Statistical analysis was performed using Prism 7 (Graphpad Software).

## Results

### MiR-148a Reduced (P)RR Expression and Protein Abundance

Using TargetScan (http://www.targetscan.org/), we performed *in silicon* analysis and found that there were two putative binding sites for miR-148a in the 3’-UTR of the (P)RR (**Figure 1A**). We thus asked whether miR-148a could regulate (P)RR expression and functions. Firstly, we tested the effects of miR-148a on *(P)RR* expression in human hepatic cells. Transfecting HepG2 and Huh7 cells with miR-148a mimics effectively reduced (P)RR transcript levels by ~60-70% (**Figure 1B**), and consequently (P)RR protein abundance (**Figure 1 C&D**). To confirm whether miR-148a directly regulates *(P)RR* expression, we mutated the putative bindings sites in the 3’-UTR of (P)RR (**Figure 1E**), and studied their consequences on gene expression using luciferase assay. Mutating the binding site 1, but not the binding site 2, attenuated miR-148a-induced reduction in luciferase activity (**Figure 1F**), confirming that the (P)RR is a direct target of miR-148a.

**Figure 1.**
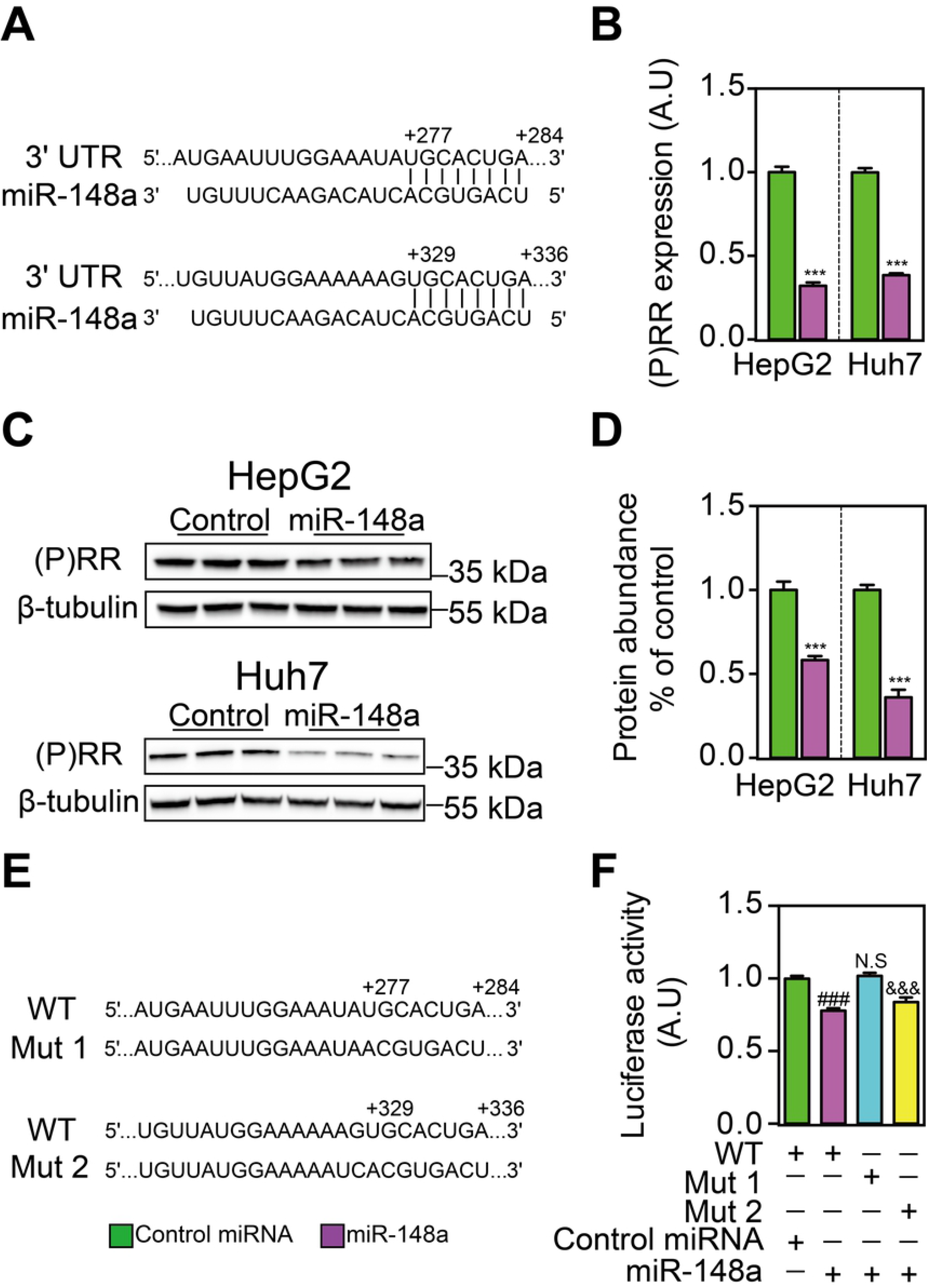
MiR-148a regulates (P)RR expression by targeting its 3’-UTR sequences. **A**. An illustration showing that there are two predicted binding sites of miR-148a on the 3’-UTR of human (P)RR. HepG2 and Huh7 cells were transfected with miR-148a or control miRNA for 48 hours, and gene expression and protein abundances were analyzed. **B.** *(P)RR* mRNA level was determined by quantitative PCR, and corrected for 36B4 level in the same sample and expressed as ratio of the control miRNA transfected. Results are from four independent experiments in triplicates. N=12; ***: p<0.001. **C**. Total cell lysates were blotted as indicated and a representative blot of 3 independent experiments in triplicates was shown. **D**. (P)RR protein abundance was quantified and normalized to the level of tubulin in the same lysates, and expressed as the relative ratio of (P)RR abundance in control miRNA transfected. N=9; ***: P<0.001. **E.** An illustration showing two constructs (Mut1 and Mut2) which are mutated for the binding site for miR-148a on the 3’-UTR of human (P)RR, comparing to wildtype (WT) sequence. **F.** HEK293T cells were transfected with luciferase reporter plasmids constructed using wildtype (WT) and mutated (Mut1 and Mut2) 3’-UTR of human (P)RR, together with either control miRNA or miR-148a. Firefly luciferase activity was measured and corrected for Renilla luciferase activity in the same sample, and expressed as ratio of WT reporter plasmid transfected samples. Results are from four independent experiments in triplicates (N=12). ###: WT+miR-148a vs. WT+control miRNA, p<0.001; N.S (not significant): Mut1+miR-148a vs. WT+miR-148a; &&&: Mut2+miR-148a vs. WT, p<0.001.

### MiR-148a Reduced LDLR Abundance and Cellular LDL uptake

Next, we tested if miR-148a could affect cellular LDL uptake as a consequence of reduced (P)RR abundance. As expected, miR-148a transfection effectively reduced cellular LDL uptake rate in HepG2 and Huh7 cells (**Figure 2 A-D**). A previous study shows that miR-148a could suppress *LDLR* expression in hepatic cells[24]. Thus, it is possible that miR-148a directly down-regulates *LDLR*, causing observed phenotype. In line with this, we found that miR-148a reduced LDLR protein abundance in HepG2 and Huh7 cells (**Figure 2 E&F**). However, LDLR transcript levels were unaltered by miR-148a transfection (**Figure 2G**), in contrast to the previous report. To further validate this observation, we generated a luciferase reporter vector by cloning the full length 3’-UTR of LDLR, and tested the effect of miR-148a on luciferase activity. Despite the presence of putative binding sites in the 3’-UTR of LDLR, miR-148a failed to reduce luciferase activity (**Figure 2H**). Moreover, transfecting the cells with siRNA targeting the 3’-UTR of LDLR potently reduced luciferase activity, validated the correctness of the reporter vector constructed (**Figure S1**). Taken together, our results suggest that miR-148a reduces cellular LDL uptake by decreasing LDLR protein abundance, without affecting LDLR transcript levels.

**Figure 2.**
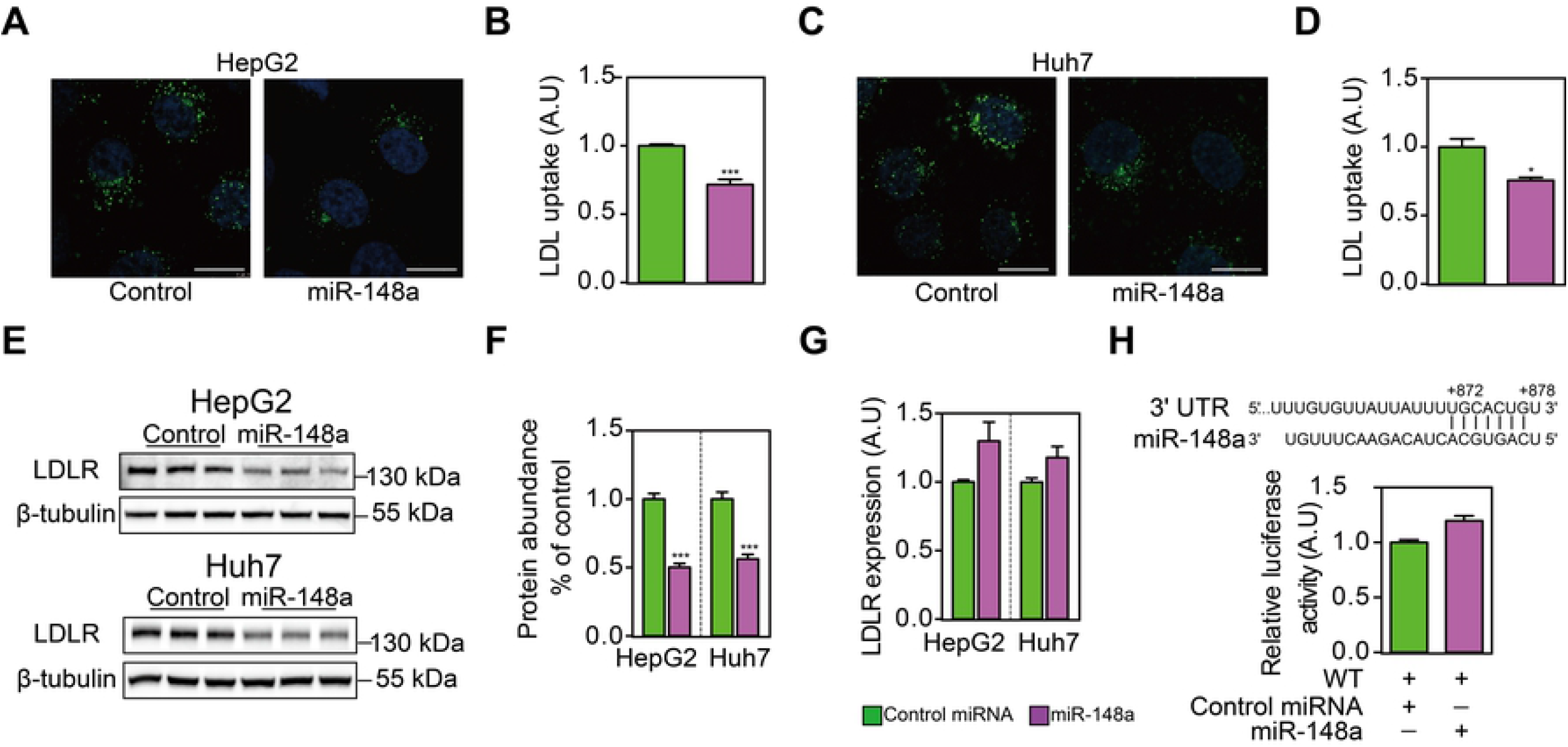
MiR-148a reduces cellular LDL uptake by reducing LDLR protein abundance but not its. HepG2 (**A & B**) and Huh7 cells (**C & D**) were transfected with control miRNA or miR-148a for 48h, and then incubated with 5 μg/mL Dylight-488-labeled LDL for 3h. **A & C**. Representative fluorescence images of cells. Nuclei are counterstained with 4’6-diaminido-2-phenylindole (blue). Scale bar: 10 μm. **B & D**. Quantitative measurement of LDL uptake in cells. Results are from four independent experiments in triplicates. N=12; *: p<0.01; ***: p<0.001. **E**. HepG2 and Huh7 cells were transfected with control miRNA or miR-148a for 48 hours. Total cell lysates were immunoblotted as indicated, and representative blots of 3 independent experiments in triplicates were shown. **F**. LDLR protein abundance was quantified and normalized to the level of tubulin in the same lysates, and expressed as the relative ratio of LDLR abundance in control miRNA transfected. N=9. ***: p<0.001. **G**. HepG2 and Huh7 cells were transfected as above, and *LDLR* mRNA levels were determined by quantitative PCR. Results are from four independent experiments in triplicates. N=12; *: p<0.05. **H**. Putative miR-148a binding sites were identified in the 3’-UTR of human LDLR, and luciferase reporter plasmid was constructed accordingly. HEK293T cells were transfected the reporter plasmid. Firefly luciferase activity was measured and corrected for Renilla luciferase activity in the same sample, and expressed as ratio of WT 3’-UTR transfected samples. Results are from four independent experiments in triplicates (N=12).

### MiR-148a Regulated LDL Uptake Via Modulating (P)RR Levels

To explore whether miR-148a regulates LDL metabolism specifically by controlling (P)RR levels, we first tested the effects of miR-148a inhibitors on (P)RR levels and cellular LDL uptake. As expected, transfecting cells with miR-148a inhibitors, but not control inhibitors, attenuated miR-148a-induced reduction in *(P)RR* transcript levels in HepG2 and Huh7 cells (**Figure 3A**). As expected, incubating cells with miR-148a or together with its inhibitors did not affect *LDLR* expression, further supporting that LDLR is not a direct target of miR-148a, at least not in the hepatic cells tested. Similar to the changes in mRNA, blocking miR-148a with its inhibitors reversed the reduction in (P)RR protein abundance caused by miR-148a transfection in hepatic cells (**Figure 3 B&C**). Antagonizing miR-148a using its inhibitors also effectively prevented miR-148a-induced reduction in LDLR protein abundance and LDL uptake (**Figure 3 D&E**), likely due to recovered (P)RR levels.

**Figure 3.**
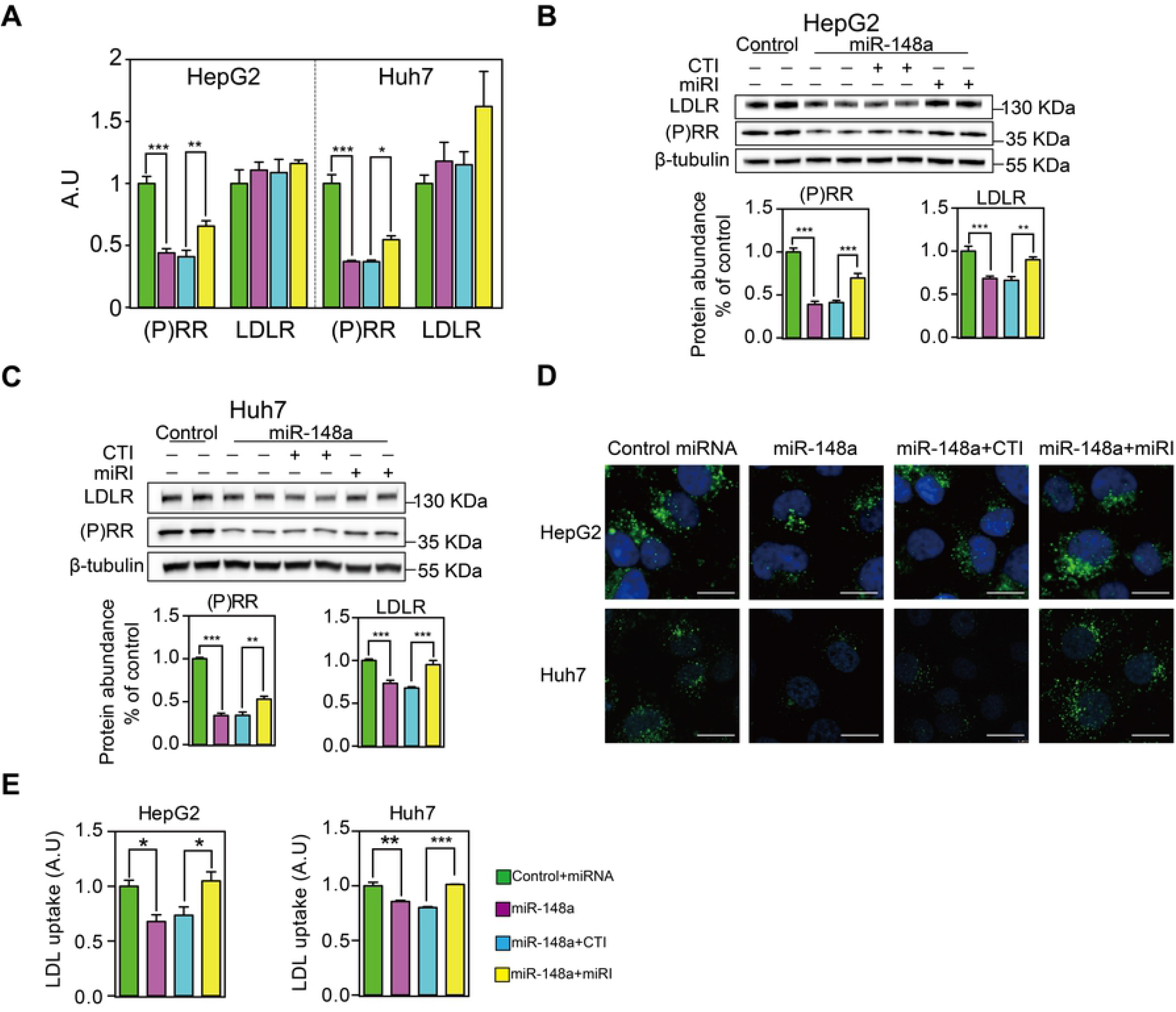
MiR-148a reduces cellular LDL uptake by specifically modulating *(P)RR* expression. HepG2 and Huh7 cells were transfected with control miRNA or miR-148a, together with control inhibitor (CTI) or miR-148a inhibitor (miRI) for 48 hours. **A**. (P)RR and LDLR mRNA levels were determined by quantitative PCR. Results are from 4 independent experiments in triplicates. N=12; **: p<0.01; ***: p<0.001. **B & C.** HepG2 (**B**) and Huh7 cells (**C**) were treated as above, and total cell lysates were immunoblotted as indicated. Representative blots of 4 independent experiments in duplicate were shown. (P)RR and LDLR protein abundance were quantified. N=8. **: p<0.01; ***: p<0.001. **D & E**. HepG2 and Huh7 cells were treated as above, and subsequently incubated with 5 μg/mL Dylight-488-labeled LDL for 3 hours. **D**. Representative fluorescence images. Scale bar: 10 μm. **E**. Quantitative measurement of LDL uptake. Results are from 3 independent experiments in triplicates. N=9; *: p<0.01; ***: p<0.001.

### Overexpressing (P)RR Reversed MiR-148a-induced LDLR Protein Reduction

To further confirm if the (P)RR is critical for miR-148a to regulate LDLR functions, we overexpressed the (P)RR in hepatic cells in the presence of miR-148a. As expected, overexpressing the (P)RR abolished miR-148a-induced reduction in LDLR protein abundance in HepG2 and Huh7 cells (**Figure 4 A&B**). As a result, miR-148a induced reduction in cellular LDL uptake in these cells was reversed by (P)RR overexpression (**Figure 4 C&D**). Collectively, our results show that miR-148a regulates LDLR abundance and LDL uptake by modulating (P)RR levels.

**Figure 4.**
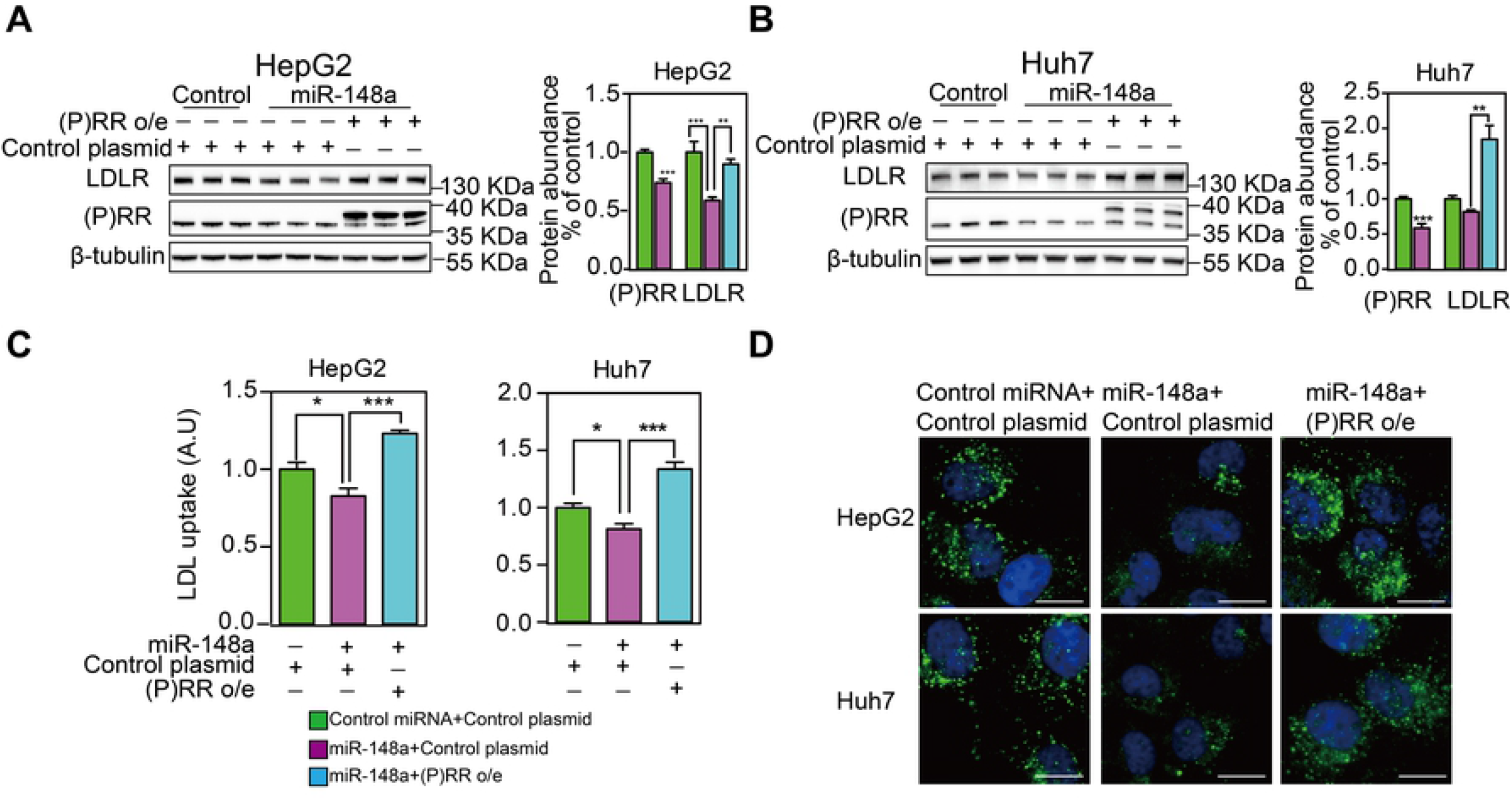
Overexpressing (P)RR antagonizes miR-148a-induced reduction in cellular LDL uptake and LDLR protein abundance. HepG2 and Huh7 cells were transfected with control miRNA or miR-148a together with control plasmid or (P)RR o/e plasmid. **A & B**. Total lysates were blotted as indicated, and representative blot of 3 independent experiments in triplicates was shown. (P)RR and LDLR protein abundance were quantified and normalized to the level of tubulin in the same lysates, and expressed as ratio of control miRNA transfected. N=9; *: p<0.05; **: p<0.01; ***: p<0.001. **C & D**. HepG2 and Huh7 cells were treated above, and subsequently incubated with 5 μg/mL Dylight488-labeled LDL for 3 hours. C. Representative fluorescence images. Scale bar, 10 μm. **D**. Quantitative measurement of LDL uptake. Results are from 3 independent experiments in triplicates. N=9; *: p<0.01; **: p<0.001.

## Discussion

In this study, we identified miR-148a as a regulator for *(P)RR* expression in hepatic cells. MiR-148a is a member of the miR-148 family, which is consisted of miR-148a, miR-148b and miR-152, sharing similarities in nucleotide sequences. Interestingly, miR-152 has previously been reported to regulate *(P)RR* expression [19]. It is likely that this family plays important role in regulating (P)RR functions. In viewing of this, we tested the effects of miR-148b on *(P)RR* expression and found miR-148b was capable of reducing (P)RR transcript levels and protein abundance, to a similar extent as miR-148a in hepatic cells (**Figure S2**).

In the current study, we found that miR-148a reduces LDLR protein abundance, and cellular LDL uptake as a consequence. Our findings are in line with a previous report showing that miR-148a regulates LDL metabolism [27]. Yet, in contrast to the previous study, we found miR-148a did not affect *LDLR* transcript levels. It is unclear why such discrepancies exist, especially the same human hepatic cell lines were used in both studies. To rule out the possibility that the cells we used were contaminated with other cells, such as HeLa, we screened 19 short tandem repeat loci and the gender determining locus to determine their identity and confirmed that HepG2 and Huh7 cells used in the current study are correct cell lines (data not shown). Furthermore, in this study, we found that reduction in LDLR protein abundance and LDL uptake induced by miR-148a were reversed by overexpressing the (P)RR, arguing that (P)RR deficiency is the underlying cause for reduced cellular LDL uptake. Decreased (P)RR abundance will also cause reduction in protein abundance of sortilin-1 (SORT1) [8]. SORT1 plays important roles in several cellular functions, such as neuronal survival and protein sorting [28, 29], and recently has been identified as a clearance receptor for LDL and an important determinant of circulating LDL levels [30–32]. In the current study, we found that miR-148a reduced SORT1 protein abundance but did not affect its transcript levels (**Figure S3**), in line with our previous report showing that (P)RR regulates LDLR and SORT1 protein abundance post-transcriptionally [8]. These findings together favor that miR-148a regulates LDL metabolism via modulating the (P)RR, instead of directly regulating LDLR.

Recent studies reported that miR-148a levels were increased in the liver and adipose tissues of diet-induced obese mice, and positively associated with body-mass index in human [27, 33]. It is possible that elevated miR-148a levels under such conditions contribute to dysregulation of LDL metabolism, a common disorder in obese subjects. However, the role of the (P)RR in lipid metabolism is rather complex. Inhibiting the (P)RR in the liver on one hand decreases LDL clearance but also on the other hand reduces hepatic VLDL secretion by limiting cholesterol and triglycerides synthesis in mice [16]. Moreover, inhibiting hepatic (P)RR also increases energy expenditure and thus attenuates diet-induced obesity in mice. It is thus possible that miR-148a could, by regulating (P)RR levels, play a role in regulating energy metabolism and lipid biosynthesis, in addition to regulating LDL metabolism.

Altered miR-148a levels have been observed under other pathophysiological conditions, such as in type 1 diabetes, gastrointestinal cancer, and ovarian cancer [22, 34–36]. It would be interesting to explore whether miR-148a could, by modulating (P)RR-mediated functions, such as regulating lipid metabolism, maintaining V-ATPase integrity, and activating tissue RAS, contribute to the onset and progression of such diseases. Among these diseases, it is interesting to notice that plasma miR-148a levels were elevated in LN patients and mice [37]. LN patients have higher risks for ischemic heart diseases [38, 39], which is likely caused by abnormalities in lipid metabolism. It is estimated that 30-50% of LN patients have high plasma total cholesterol and LDL levels [40–42]. It is unclear whether elevated miR-148a would contribute to hypercholesterolemia in LN patients by affecting LDL metabolism via the (P)RR. In fact, it is still a largely unsolved problem why patients with nephrotic syndrome (NS) develop hypercholesterolemia. Recent studies found that elevated hepatic IDOL and PCSK9 may contribute to NS-induced hypercholesterolemia [43], and ablation of hepatic PCSK9 attenuated NS-induced hypercholesterolemia in mice [44]. Yet, it is still unclear how renal dysfunctions in NS signals to the liver to regulate cholesterol metabolism. It is possible that miRNAs, including miR-148a, play an essential role in this process. Further studies are needed to clarify if miR-148a would contribute to hypercholesterolemia in NS.

In summary, we report that miR-148a reduces *(P)RR* expression by directly binding to the 3’-UTR of (P)RR messenger RNA in human hepatic cells. By suppressing *(P)RR* expression, miR-148a reduces LDLR protein abundance and consequently LDL metabolism. Our study highlights that miR-148a is a regulator for (P)RR expression and functions.

## Acknowledgement

This work is supported by National Natural Science Foundation of China (grant no. 81870605, 81800383, 81500667) and Shenzhen Municipal Science and Technology Innovation Council (grant no. JCYJ20160307160819191).

## References

1. Thedrez A, Blom DJ, Ramin-Mangata S, Blanchard V, Croyal M, Chemello K, et al. Homozygous Familial Hypercholesterolemia Patients With Identical Mutations Variably Express the LDLR (Low-Density Lipoprotein Receptor): Implications for the Efficacy of Evolocumab. Arterioscler Thromb Vasc Biol. 2018;38(3):592–8. Epub 2017/12/30. doi: 10.1161/ATVBAHA.117.310217. PubMed PMID: 29284604; PubMed Central PMCID: PMCPMC5823753.

2. Brown MS, Goldstein JL. The SREBP pathway: regulation of cholesterol metabolism by proteolysis of a membrane-bound transcription factor. Cell. 1997;89(3):331–40. Epub 1997/05/02. doi: 10.1016/s0092-8674(00)80213-5. PubMed PMID: 9150132.

3. Maxwell KN, Fisher EA, Breslow JL. Overexpression of PCSK9 accelerates the degradation of the LDLR in a post-endoplasmic reticulum compartment. Proc Natl Acad Sci U S A. 2005;102(6):2069–74. Epub 2005/01/29. doi: 10.1073/pnas.0409736102. PubMed PMID: 15677715; PubMed Central PMCID: PMCPMC546019.

4. Sorrentino V, Zelcer N. Post-transcriptional regulation of lipoprotein receptors by the E3-ubiquitin ligase inducible degrader of the low-density lipoprotein receptor. Curr Opin Lipidol. 2012;23(3):213–9. Epub 2012/04/19. doi: 10.1097/MOL.0b013e3283532947. PubMed PMID: 22510808.

5. Zelcer N, Hong C, Boyadjian R, Tontonoz P. LXR regulates cholesterol uptake through Idol-dependent ubiquitination of the LDL receptor. Science. 2009;325(5936):100–4. Epub 2009/06/13. doi: 10.1126/science.1168974. PubMed PMID: 19520913; PubMed Central PMCID: PMCPMC2777523.

6. Cohen J, Pertsemlidis A, Kotowski IK, Graham R, Garcia CK, Hobbs HH. Low LDL cholesterol in individuals of African descent resulting from frequent nonsense mutations in PCSK9. Nat Genet. 2005;37(2):161–5. Epub 2005/01/18. doi: 10.1038/ng1509. PubMed PMID: 15654334.

7. Horton JD, Cohen JC, Hobbs HH. Molecular biology of PCSK9: its role in LDL metabolism. Trends Biochem Sci. 2007;32(2):71–7. Epub 2007/01/12. doi: 10.1016/j.tibs.2006.12.008. PubMed PMID: 17215125; PubMed Central PMCID: PMCPMC2711871.

8. Lu X, Meima ME, Nelson JK, Sorrentino V, Loregger A, Scheij S, et al. Identification of the (Pro)renin Receptor as a Novel Regulator of Low-Density Lipoprotein Metabolism. Circ Res. 2016;118(2):222–9. Epub 2015/11/20. doi: 10.1161/CIRCRESAHA.115.306799. PubMed PMID: 26582775.

9. Feldt S, Batenburg WW, Mazak I, Maschke U, Wellner M, Kvakan H, et al. Prorenin and renin-induced extracellular signal-regulated kinase 1/2 activation in monocytes is not blocked by aliskiren or the handle-region peptide. Hypertension. 2008;51(3):682–8. Epub 2008/01/24. doi: 10.1161/HYPERTENSIONAHA.107.101444. PubMed PMID: 18212269.

10. Huang Y, Noble NA, Zhang J, Xu C, Border WA. Renin-stimulated TGF-beta1 expression is regulated by a mitogen-activated protein kinase in mesangial cells. Kidney Int. 2007;72(1):45–52. Epub 2007/03/31. doi: 10.1038/sj.ki.5002243. PubMed PMID: 17396111.

11. Liu Y, Zhang S, Su D, Liu J, Cheng Y, Zou L, et al. Inhibiting (pro)renin receptor-mediated p38 MAPK signaling decreases hypoxia/reoxygenation-induced apoptosis in H9c2 cells. Mol Cell Biochem. 2015;403(1-2):267–76. Epub 2015/02/26. doi: 10.1007/s11010-015-2356-8. PubMed PMID: 25711402.

12. Cruciat CM, Ohkawara B, Acebron SP, Karaulanov E, Reinhard C, Ingelfinger D, et al. Requirement of prorenin receptor and vacuolar H+-ATPase-mediated acidification for Wnt signaling. Science. 2010;327(5964):459–63. Epub 2010/01/23. doi: 10.1126/science.1179802. PubMed PMID: 20093472.

13. Hermle T, Saltukoglu D, Grunewald J, Walz G, Simons M. Regulation of Frizzled-dependent planar polarity signaling by a V-ATPase subunit. Curr Biol. 2010;20(14):1269–76. Epub 2010/06/29. doi: 10.1016/j.cub.2010.05.057. PubMed PMID: 20579879.

14. Kinouchi K, Ichihara A, Sano M, Sun-Wada GH, Wada Y, Kurauchi-Mito A, et al. The (pro)renin receptor/ATP6AP2 is essential for vacuolar H+-ATPase assembly in murine cardiomyocytes. Circ Res. 2010;107(1):30–4. Epub 2010/06/24. doi: 10.1161/CIRCRESAHA.110.224667. PubMed PMID: 20570919.

15. Li Z, Zhou L, Wang Y, Miao J, Hong X, Hou FF, et al. (Pro)renin Receptor Is an Amplifier of Wnt/beta-Catenin Signaling in Kidney Injury and Fibrosis. J Am Soc Nephrol. 2017;28(8):2393–408. Epub 2017/03/09. doi: 10.1681/ASN.2016070811. PubMed PMID: 28270411; PubMed Central PMCID: PMCPMC5533230.

16. Ren L, Sun Y, Lu H, Ye D, Han L, Wang N, et al. (Pro)renin Receptor Inhibition Reprograms Hepatic Lipid Metabolism and Protects Mice From Diet-Induced Obesity and Hepatosteatosis. Circ Res. 2018;122(5):730–41. Epub 2018/01/06. doi: 10.1161/CIRCRESAHA.117.312422. PubMed PMID: 29301853; PubMed Central PMCID: PMCPMC6309662.

17. Cheng H, Fan X, Moeckel GW, Harris RC. Podocyte COX-2 exacerbates diabetic nephropathy by increasing podocyte (pro)renin receptor expression. J Am Soc Nephrol. 2011;22(7):1240–51. Epub 2011/07/09. doi: 10.1681/ASN.2010111149. PubMed PMID: 21737546; PubMed Central PMCID: PMCPMC3137572.

18. Schefe JH, Menk M, Reinemund J, Effertz K, Hobbs RM, Pandolfi PP, et al. A novel signal transduction cascade involving direct physical interaction of the renin/prorenin receptor with the transcription factor promyelocytic zinc finger protein. Circ Res. 2006;99(12):1355–66. Epub 2006/11/04. doi: 10.1161/01.RES.0000251700.00994.0d. PubMed PMID: 17082479.

19. Haque R, Hur EH, Farrell AN, Iuvone PM, Howell JC. MicroRNA-152 represses VEGF and TGFbeta1 expressions through post-transcriptional inhibition of (Pro)renin receptor in human retinal endothelial cells. Mol Vis. 2015;21:224–35. Epub 2015/03/25. PubMed PMID: 25802486; PubMed Central PMCID: PMCPMC4358229.

20. Filipowicz W, Bhattacharyya SN, Sonenberg N. Mechanisms of post-transcriptional regulation by microRNAs: are the answers in sight? Nat Rev Genet. 2008;9(2):102–14. Epub 2008/01/17. doi: 10.1038/nrg2290. PubMed PMID: 18197166.

21. Batkai S, Thum T. MicroRNAs in hypertension: mechanisms and therapeutic targets. Curr Hypertens Rep. 2012;14(1):79–87. Epub 2011/11/05. doi: 10.1007/s11906-011-0235-6. PubMed PMID: 22052337.

22. Horie T, Baba O, Kuwabara Y, Chujo Y, Watanabe S, Kinoshita M, et al. MicroRNA-33 deficiency reduces the progression of atherosclerotic plaque in ApoE-/-mice. J Am Heart Assoc. 2012;1(6):e003376. Epub 2013/01/15. doi: 10.1161/JAHA.112.003376. PubMed PMID: 23316322; PubMed Central PMCID: PMCPMC3540673.

23. Ruan Q, Wang T, Kameswaran V, Wei Q, Johnson DS, Matschinsky F, et al. The microRNA-21-PDCD4 axis prevents type 1 diabetes by blocking pancreatic beta cell death. Proc Natl Acad Sci U S A. 2011;108(29):12030–5. Epub 2011/07/07. doi: 10.1073/pnas.1101450108. PubMed PMID: 21730150; PubMed Central PMCID: PMCPMC3141944.

24. Goedeke L, Rotllan N, Canfran-Duque A, Aranda JF, Ramirez CM, Araldi E, et al. MicroRNA-148a regulates LDL receptor and ABCA1 expression to control circulating lipoprotein levels. Nat Med. 2015;21(11):1280–9. Epub 2015/10/06. doi: 10.1038/nm.3949. PubMed PMID: 26437365; PubMed Central PMCID: PMCPMC4711995.

25. Kleinveld HA, Duif PF, Pekelharing HL, van Rijn HJ. Oxidation of lipoprotein(a) and low density lipoprotein containing density gradient ultracentrifugation fractions. Biochim Biophys Acta. 1996;1303(1):15–21. Epub 1996/09/06. doi: 10.1016/0005-2760(96)00055-0. PubMed PMID: 8816848.

26. Zhang J, Noble NA, Border WA, Owens RT, Huang Y. Receptor-dependent prorenin activation and induction of PAI-1 expression in vascular smooth muscle cells. Am J Physiol Endocrinol Metab. 2008;295(4):E810–9. Epub 2008/07/31. doi: 10.1152/ajpendo.90264.2008. PubMed PMID: 18664599; PubMed Central PMCID: PMCPMC2575903.

27. Wagschal A, Najafi-Shoushtari SH, Wang L, Goedeke L, Sinha S, deLemos AS, et al. Genome-wide identification of microRNAs regulating cholesterol and triglyceride homeostasis. Nat Med. 2015;21(11):1290–7. Epub 2015/10/27. doi: 10.1038/nm.3980. PubMed PMID: 26501192; PubMed Central PMCID: PMCPMC4993048.

28. Campbell C, Beug S, Nickerson PE, Peng J, Mazerolle C, Bassett EA, et al. Sortilin regulates sorting and secretion of Sonic hedgehog. J Cell Sci. 2016;129(20):3832–44. Epub 2016/09/17. doi: 10.1242/jcs.183541. PubMed PMID: 27632999.

29. Jansen P, Giehl K, Nyengaard JR, Teng K, Lioubinski O, Sjoegaard SS, et al. Roles for the pro-neurotrophin receptor sortilin in neuronal development, aging and brain injury. Nat Neurosci. 2007;10(11):1449–57. Epub 2007/10/16. doi: 10.1038/nn2000. PubMed PMID: 17934455.

30. Linsel-Nitschke P, Heeren J, Aherrahrou Z, Bruse P, Gieger C, Illig T, et al. Genetic variation at chromosome 1p13.3 affects sortilin mRNA expression, cellular LDL-uptake and serum LDL levels which translates to the risk of coronary artery disease. Atherosclerosis. 2010;208(1):183–9. Epub 2009/08/08. doi: 10.1016/j.atherosclerosis.2009.06.034. PubMed PMID: 19660754.

31. Musunuru K, Strong A, Frank-Kamenetsky M, Lee NE, Ahfeldt T, Sachs KV, et al. From noncoding variant to phenotype via SORT1 at the 1p13 cholesterol locus. Nature. 2010;466(7307):714–9. doi: 10.1038/nature09266.

32. Strong A, Ding Q, Edmondson AC, Millar JS, Sachs KV, Li X, et al. Hepatic sortilin regulates both apolipoprotein B secretion and LDL catabolism. Journal of Clinical Investigation. 2012;122(8):2807–16. doi: 10.1172/jci63563.

33. Shi C, Zhang M, Tong M, Yang L, Pang L, Chen L, et al. miR-148a is Associated with Obesity and Modulates Adipocyte Differentiation of Mesenchymal Stem Cells through Wnt Signaling. Sci Rep. 2015;5:9930. Epub 2015/05/23. doi: 10.1038/srep09930. PubMed PMID: 26001136; PubMed Central PMCID: PMCPMC4441322.

34. Chen Y, Song Y, Wang Z, Yue Z, Xu H, Xing C, et al. Altered expression of MiR-148a and MiR-152 in gastrointestinal cancers and its clinical significance. J Gastrointest Surg. 2010;14(7):1170–9. Epub 2010/04/28. doi: 10.1007/s11605-010-1202-2. PubMed PMID: 20422307.

35. Grieco GE, Cataldo D, Ceccarelli E, Nigi L, Catalano G, Brusco N, et al. Serum Levels of miR-148a and miR-21-5p Are Increased in Type 1 Diabetic Patients and Correlated with Markers of Bone Strength and Metabolism. Noncoding RNA. 2018;4(4). Epub 2018/11/30. doi: 10.3390/ncrna4040037. PubMed PMID: 30486455; PubMed Central PMCID: PMCPMC6315714.

36. Zhou X, Zhao F, Wang ZN, Song YX, Chang H, Chiang Y, et al. Altered expression of miR-152 and miR-148a in ovarian cancer is related to cell proliferation. Oncol Rep. 2012;27(2):447–54. Epub 2011/10/06. doi: 10.3892/or.2011.1482. PubMed PMID: 21971665.

37. Qingjuan L, Xiaojuan F, Wei Z, Chao W, Pengpeng K, Hongbo L, et al. miR-148a-3p overexpression contributes to glomerular cell proliferation by targeting PTEN in lupus nephritis. Am J Physiol Cell Physiol. 2016;310(6):C470–8. Epub 2016/01/23. doi: 10.1152/ajpcell.00129.2015. PubMed PMID: 26791485.

38. Sinicato NA, da Silva Cardoso PA, Appenzeller S. Risk factors in cardiovascular disease in systemic lupus erythematosus. Curr Cardiol Rev. 2013;9(1):15–9. Epub 2013/03/08. PubMed PMID: 23463953; PubMed Central PMCID: PMCPMC3584302.

39. Sui M, Jia X, Yu C, Guo X, Liu X, Ji Y, et al. Relationship between hypoalbuminemia, hyperlipidemia and renal severity in patients with lupus nephritis: a prospective study. Cent Eur J Immunol. 2014;39(2):243–52. Epub 2014/01/01. doi: 10.5114/ceji.2014.43730. PubMed PMID: 26155131; PubMed Central PMCID: PMCPMC4440014.

40. Sajjad S, Farman S, Saeed MA, Ahmad NM, Butt BA. Frequency of Dyslipidemia in patients with Lupus Nephritis. Pak J Med Sci. 2017;33(2):358–62. Epub 2017/05/20. doi: 10.12669/pjms.332.12410. PubMed PMID: 28523037; PubMed Central PMCID: PMCPMC5432704.

41. Ali Abdalla M, Mostafa El Desouky S, Sayed Ahmed A. Clinical significance of lipid profile in systemic lupus erythematosus patients: Relation to disease activity and therapeutic potential of drugs. The Egyptian Rheumatologist. 2017;39(2):93–8. doi: 10.1016/j.ejr.2016.08.004.

42. Faurschou M, Mellemkjaer L, Starklint H, Kamper AL, Tarp U, Voss A, et al. High risk of ischemic heart disease in patients with lupus nephritis. J Rheumatol. 2011;38(11):2400–5. Epub 2011/09/03. doi: 10.3899/jrheum.110329. PubMed PMID: 21885497.

43. Liu S, Vaziri ND. Role of PCSK9 and IDOL in the pathogenesis of acquired LDL receptor deficiency and hypercholesterolemia in nephrotic syndrome. Nephrol Dial Transplant. 2014;29(3):538–43. Epub 2013/10/30. doi: 10.1093/ndt/gft439. PubMed PMID: 24166456.

44. Haas ME, Levenson AE, Sun X, Liao WH, Rutkowski JM, de Ferranti SD, et al. The Role of Proprotein Convertase Subtilisin/Kexin Type 9 in Nephrotic Syndrome-Associated Hypercholesterolemia. Circulation. 2016;134(1):61–72. Epub 2016/07/01. doi: 10.1161/CIRCULATIONAHA.115.020912. PubMed PMID: 27358438; PubMed Central PMCID: PMCPMC5345853.

